# Alteration of mucins in the submandibular gland during aging in mice

**DOI:** 10.1101/2020.08.12.247478

**Authors:** Akihiko Kameyama, Wai Wai Thet Tin, Risa Nishijima, Kimi Yamakoshi

## Abstract

**Objective:** Mucins are large glycosylated glycoproteins that are produced in the salivary glands, and their changes may contribute to the development of xerostomia due to aging and the accompanying deterioration of oral hygiene. This study aimed to characterize the changes in the mucins produced in submandibular gland (SMG) during the aging process.

**Methods:** SMG mucins derived from mice of each age were separated using supported molecular matrix electrophoresis (SMME). Subsequently, the membranes were stained with AB or blotted with MAL-II lectin. The SMME membranes stained with AB were subjected to densitometric analysis and glycan analysis. The detailed structures of O-glycan were investigated by MS/MS spectra.

**Results:** The SMG of mice secreted three mucins with different glycan profiles: age-specific mucin, youth-specific mucin, and a mucin expressed throughout life, and the expression patterns of these mucins change during aging. Additionally, age-specific mucin began to be detected at about 12 months of age. A mucin expressed throughout life and age-specific mucin had the same mass of major glycans but different structures.

Furthermore, the proportion of mucin glycan species expressed throughout life changed during the aging process, and aging tended to decrease the proportion of fucosylated glycans and increase the proportion of sialoglycans.

**Conclusion:** There are three secretory mucins with different glycan profiles in the SMG of mice, and their expression patterns change according to the period of the aging process. The proportion of glycan species of mucin expressed throughout life also changes during the aging process.

**Highlights:**

- Three secreted mucins with different glycan profiles are detected in SMG using SMME.
- Each mucin has its own peak production age during the aging process.
- Age-specific mucin begins to be detected at about 12 months of age.
- Same mass of major glycans but different structures in mucins expressed throughout life vs. age-specific mucins.
- Proportion of mucin glycan species expressed throughout life changes during aging.

## Introduction

Saliva is indispensable for the maintenance of oral health (Dodds, Johnson, & Yeh, 2005; Edgar, 1992). In many cases, however, the elderly suffer from poor oral health. This is mainly due to xerostomia, which is defined as the subjective perception of dry mouth typically associated with reduced salivary flow and increased salivary viscosity in many elderly people (Hopcraft & Tan, 2010; Osterberg, Landahl, & Hedegard, 1984; Sreebny, 2000; Zussman, Yarin, & Nagler, 2007). Xerostomia and its associated deterioration in oral hygiene among the elderly not only predisposes this population to oral diseases such as dental caries and periodontal disease (Abdellatif & Burt, 1987; Gilbert, Heft, & Duncan, 1993) but may also have systemic adverse effects such as an increased risk of aspiration pneumonia (Sjogren, Nilsson, Forsell, Johansson, & Hoogstraate, 2008; van der Maarel-Wierink, Vanobbergen, Bronkhorst, Schols, & de Baat, 2013), poor nutritional status, and impaired cognitive function (Harding, Gonder, Robinson, Crean, & Singhrao, 2017; Rhodus & Brown, 1990). A better understanding of the factors that cause xerostomia is likely to facilitate the development of new strategies for its prevention and treatment.

Xerostomia in the elderly may involve critical changes in mucins, which are produced in the salivary glands and are major organic components of salivary mucus. Mucins are large mass glycoproteins in which the core protein has been modified with O-linked glycans accounting for 50%–80% of the mass. Mucins bind to microorganisms in the microbiota via interaction of their mucin glycans with adhesion molecules on the surface of the microorganism. Aggregates of mucin in the saliva cleanse microorganisms from the oral cavity, while mucins that cover the surfaces of the teeth and mucous membranes promote the colonization of microorganisms on these surfaces (Humphrey & Williamson, 2001; Wu, Csako, & Herp, 1994). Thus, the interaction of these opposing forces contributes to the maintenance of oral hygiene through selective regulation of the adhesion of microorganisms to oral tissue surfaces by salivary mucins and the control of bacterial colonization. However, mucin oligomerization scales and glycosylation structures are greatly varied, and the expression and glycan structure of mucins are easily altered by environment (Hollingsworth & Swanson, 2004; Robbe et al., 2003; Voynow, Gendler, & Rose, 2006). Alterations in mucin glycan structure affect not only physical properties such as gel formation by mucin oligomerization but also affect many complex microflora-mucin interactions (Derrien et al., 2010; Robbe et al., 2003). Considering that salivary viscosity increases and oral hygiene worsens in elderly people with xerostomia, it is possible that changes in the physiochemical composition of mucin molecules may alter the functions of mucin for microflora during the aging process. However, because mucins are difficult to analyze due to their large mass and complex biochemical composition, changes in SG mucin during the aging process have not yet been characterized.

Using supported molecular matrix electrophoresis (SMME), a method developed to separate high molecular weight glycoproteins (Matsuno, Saito, Gotoh, Narimatsu, & Kameyama, 2009), we reported that aging causes the production of new mucin-like molecules in the SMG, one of the SGs of mice (Iida et al., 2019; Kameyama, Yamakoshi, & Watanabe, 2019). Unlike polyacrylamide and agarose gel electrophoresis, this method electrophoreses mucins regardless of their molecular weights. In this study, to examine the SMG mucin profile during the aging process, we performed SMME with a slight modification from the previously described mucin enrichment method, visualized the mucin on the electrophoresed SMME membrane by staining glycosaminoglycans and acidic mucins such as sialomucin and sulfomucin, and subsequently analyzed the glycan structure of these mucins by mass spectrometry (MS). We thereby found that the mouse SMG produces three secreted mucins with different glycan profiles: age-specific mucin, youth-specific mucin, and a mucin expressed throughout life, and that the expression patterns of these mucins change during aging. Youth-specific mucin is detected from youth to about 12 months of age, while age-specific mucin is detected from about 12 months of age. Furthermore, we have revealed that among the three mucins, the proportion of mucin glycan species expressed throughout life changes during the aging process, and that aging tends to decrease the proportion of fucosylated glycans and increase the proportion of sialoglycans.

## Materials and Methods

### Mice

For SMME and glycan analysis, adult C57BL/6N mice from 3 to 24 months of age were obtained from the Experimental Animal Facility at the National Center for Geriatrics and Gerontology (Obu, Aichi, Japan). Male mice were used for all experiments. Mice were sacrificed, and the SMGs were immediately isolated. All mice were housed in a pathogen-free barrier environment during the study. All animal studies were performed in adherence to the Institutional Animal Care approval number 31-29.

### Enrichment of mucins from SMG

Mucins were enriched from SMG tissue specimens according to our previous report (21). Briefly, acetone powders prepared from the SMG specimens (approximately 50 mg) were suspended in PBS (pH 7.4, 500 µL) and centrifuged at 12,000 ×*g* for 10 min at 4°C. Then the supernatants were collected in new tubes and mixed with a saturated aqueous solution of calcium acetate in 5:1 ratio. The solutions were mixed with three volumes of ethanol, kept at −80°C for 1 h, and then centrifuged at 15,000 ×*g* for 10 min at 4°C. The supernatants were discarded, and the precipitates were suspended in 2 M urea in PBS (pH 7.4, 100 µL) and centrifuged at 15,000 ×*g* for 10 min at 4°C.

Supernatants were collected in new tubes, and precipitation using ethanol was repeated as described above. The resulting precipitates were dissolved in 50 µL of 8 M urea in PBS (pH 7.4) and were stored at 4°C until use.

### Supported molecular matrix electrophoresis

SMME membranes were prepared as described in a previous report (Matsuno et al., 2009). Polyvinylidene difluoride (PVDF) membranes (Immobilon-P, pore size 0.45 μm, Millipore Corp.) were cut in a rectangular shape (6 cm × 5 cm) and immersed in 0.1% hydrophilic polymer solution with gentle shaking for 2 h. The hydrophilic polymers used for glycan analysis and lectin blotting were polyvinyl alcohol (PVA: average MW, 22,000; Wako Pure Chemical, Osaka, Japan) and 1:3 mixture of PVA and polyvinylpyrrolidone (PVP: average MW, 40,000; Sigma-Aldrich, St. Louis, MO), respectively. The prepared SMME membranes were equilibrated using a running buffer of 0.1 M pyridine-formic acid (pH 4.0) for 30 min with gentle shaking; subsequently, the SMME membranes (separation length, 6 cm) were placed in a membrane electrophoresis chamber (EPC105AA; Advantec, Tokyo, Japan). Next, 1 µL of the sample solution was streaked onto the membrane (5 mm wide and 1.2 cm from the bottom of the membrane) and subjected to a constant current at 1.0 mA/cm for 30 min. For alcian blue (AB) staining, the electrophoresed membranes were immersed in 30% acetic acid in methanol for 30 min immediately after electrophoresis and subsequently incubated in the AB dye solution (pH 4.0) with gentle shaking for 30 min. The stained membrane was washed with methanol for a few minutes to remove the background color.

### Densitometric analysis

The stained membranes were dried at 45°C for 30 min on a heating block and then were rewetted with ethylene glycol to make the membranes transparent. The membranes were imaged using the ChemiDoc XRS imaging system (Bio-Rad, Hercules, CA, USA). The obtained images were densitometrically analyzed using Image J 1.52a (NIH, http://imagej.nih.gov/ij).

### Glycan release and permethylation

Mucin bands stained with AB were excised and subjected to reductive β-elimination, as described previously (Matsuno et al., 2009). Briefly, the excised bands were incubated in 40 µL of 500 mM NaBH_4_ in 50 mM NaOH, incubated at 45°C for 16 h, and subsequently quenched by adding 8 µL of glacial acetic acid. The solutions were desalinated using cation-exchange solid phase extraction cartridge (Oasis MCX, 60 mg, Waters Corp., Milford, MA) and concentrated and co-evaporated using 1% acetic acid in methanol to remove boric acid. The resulting residues were permethylated using iodomethane and NaOH, desalinated, and dried as previously reported (Matsuno et al., 2009).

### Mass spectrometry

MS spectra were acquired using a matrix-assisted laser desorption/ionization time-of-flight (MALDI-TOF) mass spectrometer (UltraFlex; Bruker Daltonik, Bemen, Germany). Ions were generated by a pulsed 337-nm nitrogen laser and accelerated to 20 kV. All spectra were obtained in the positive reflectron mode and analyzed using FlexAnalysis 2.0 software (Bruker Daltonik). Glycan signals were assigned using GlycoMod (https://web.expasy.org/glycomod/). Tandem mass spectra were acquired using a MALDI-quadrupole ion trap (QIT)-TOF mass spectrometer (AXIMA-QIT; Shimadzu Corp., Kyoto, Japan) using a positive ion mode. Ions were generated using a 337-nm nitrogen laser and argon gas was used as a collision gas for collision-induced dissociation. For sample preparation, 0.5 µL of 2,5-dihydroxybenzoic acid (DHB) solution (10 mg/mL in 30% ethanol) was spotted onto a MALDI target plate and dried. The permethylated glycans were dissolved in 50% acetonitrile, and 0.5-µL aliquots of the solution were spotted onto the dried DHB crystal on the target plate and air dried.

### Lectin blotting with MAL-II lectin

The SMME membrane was stained with lectin in a manner similar to the previously described immunostaining [22,23]. Briefly, the membranes were transferred in acetone immediately after electrophoresis, and gently shaken for 30 min. After drying the membranes in a fume hood, heat treatment (150°C for 5 min) was performed using a thin-layer chromatography thermal blotter (ATTO Corp., Tokyo, Japan). The membranes were blocked with 5% BSA and then incubated with biotin-labeled Maackia amurensis lectin II (MAL-II biotin conjugate, Vector Laboratories, Inc., Burlingame, CA) for 1 h at room temperature. After washing with PBS-T, the membranes were incubated with horseradish peroxidase-labeled streptavidin, washed, and visualized using a chemiluminescence reagent (Western Lightning Plus-ECL, PerkinElmer, Boston, MA).

### Statistical analysis

The sample size (n) and data of mice of each age are described in Figure 1B, Figure 2 and Figure 4. Statistical significance was determined by one-way analysis of variance using GraphPad Prism 8.4.0 software (GraphPad Software Corp., CA, USA). Values of p < 0.05 were considered statistically significant.

**Figure 1.**
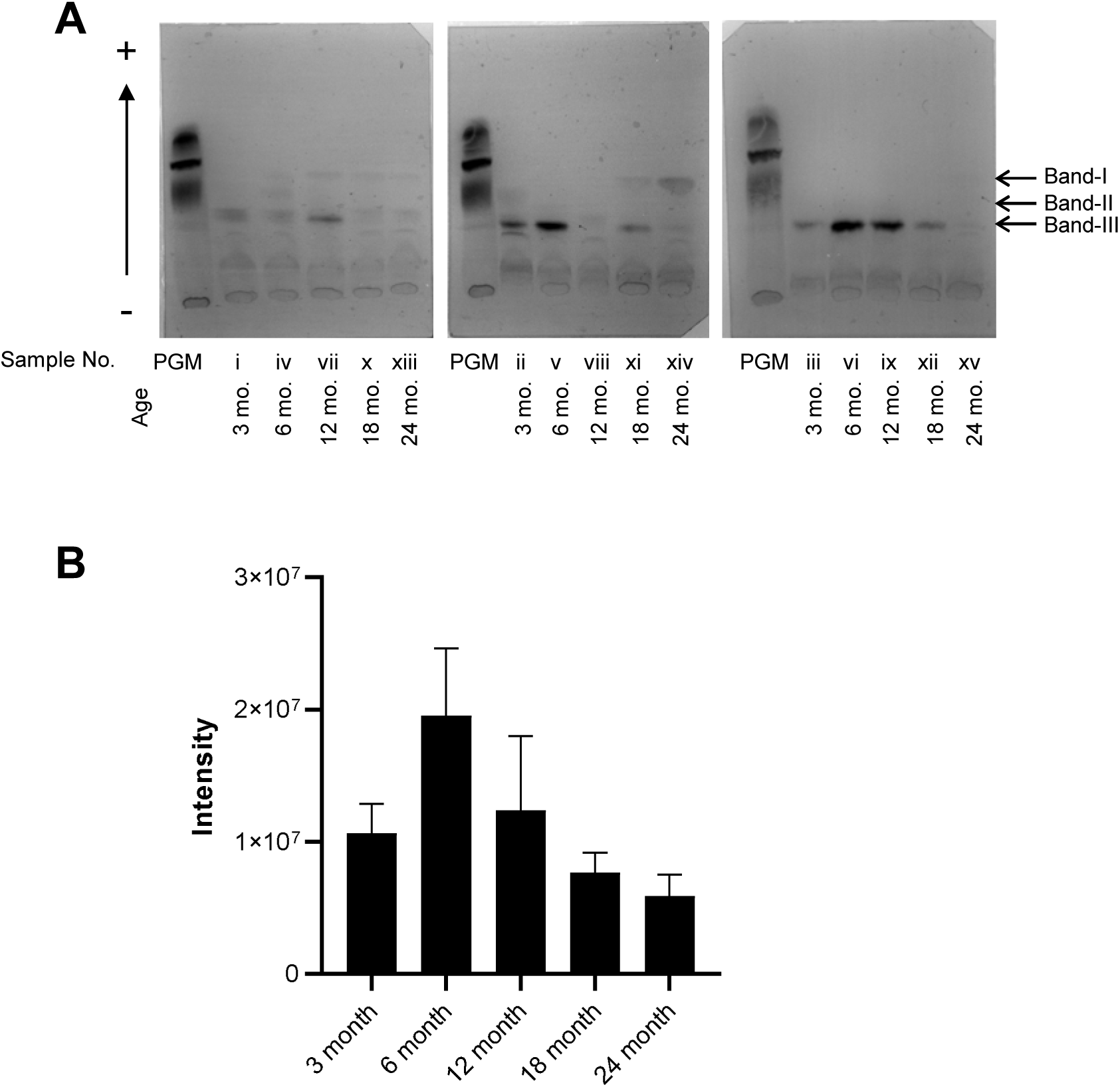
Supported molecular matrix electrophoresis analyses of submandibular gland mucin during aging in mice. (A) Comparison of mouse submandibular gland (SMG) mucin at different ages (3, 6, 12, 18, 24 months) separated by supported molecular matrix electrophoresis (SMME). Three mice were used for each age. SMME membranes are all from different individuals. Reference mucin: porcine gastric mucin (PGM). SMME membranes were stained with Alcian blue (pH 4.0). (B) Densitometric analysis of the mucin bands detected in (A). Bars represent the average of the sum of the intensity of band (I + II + III) at each age. Data are presented as means ± standard error (s.e.); *n* = 3.

**Figure 2.**
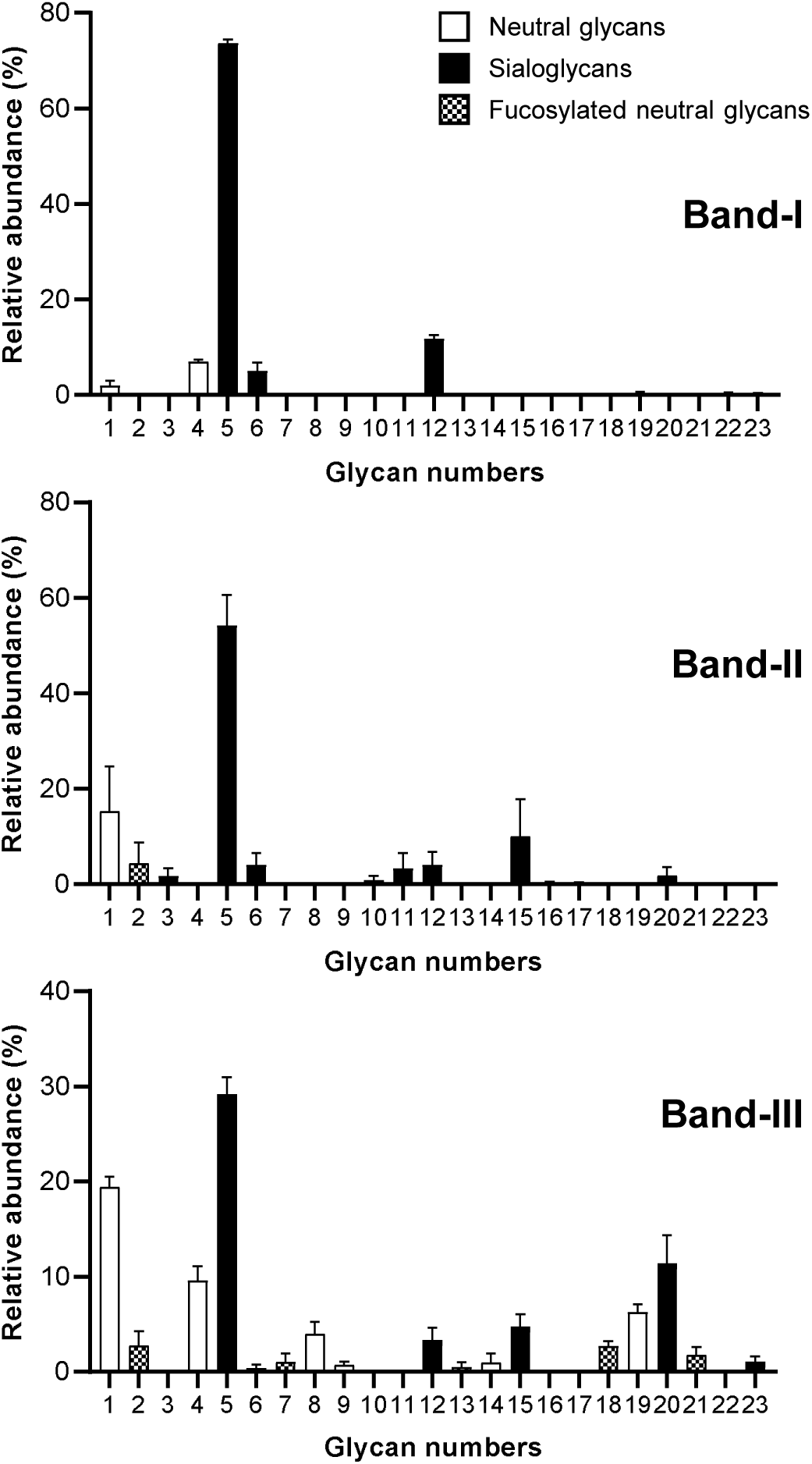
Glycan profiles of three mucins in the mouse SMG. Average glycan profiles of the three bands (Band-I, II, and III). Bars represent the relative abundance of each glycan to the whole signal intensity of permethylated glycans observed in MALDI-MS spectra. Glycan composition and *m/z* value of each glycan are summarized in Table 1. Data are presented as means ± s.e.; band-I, *n* = 6 (sample no. iii, iv, v, ix, x, xv); band-II, *n* = 5 (sample no. i, ii, iii, vi, xii); band-III, *n* = 15 (sample no. i–xv) in Figure 1A.

**Figure 3.**
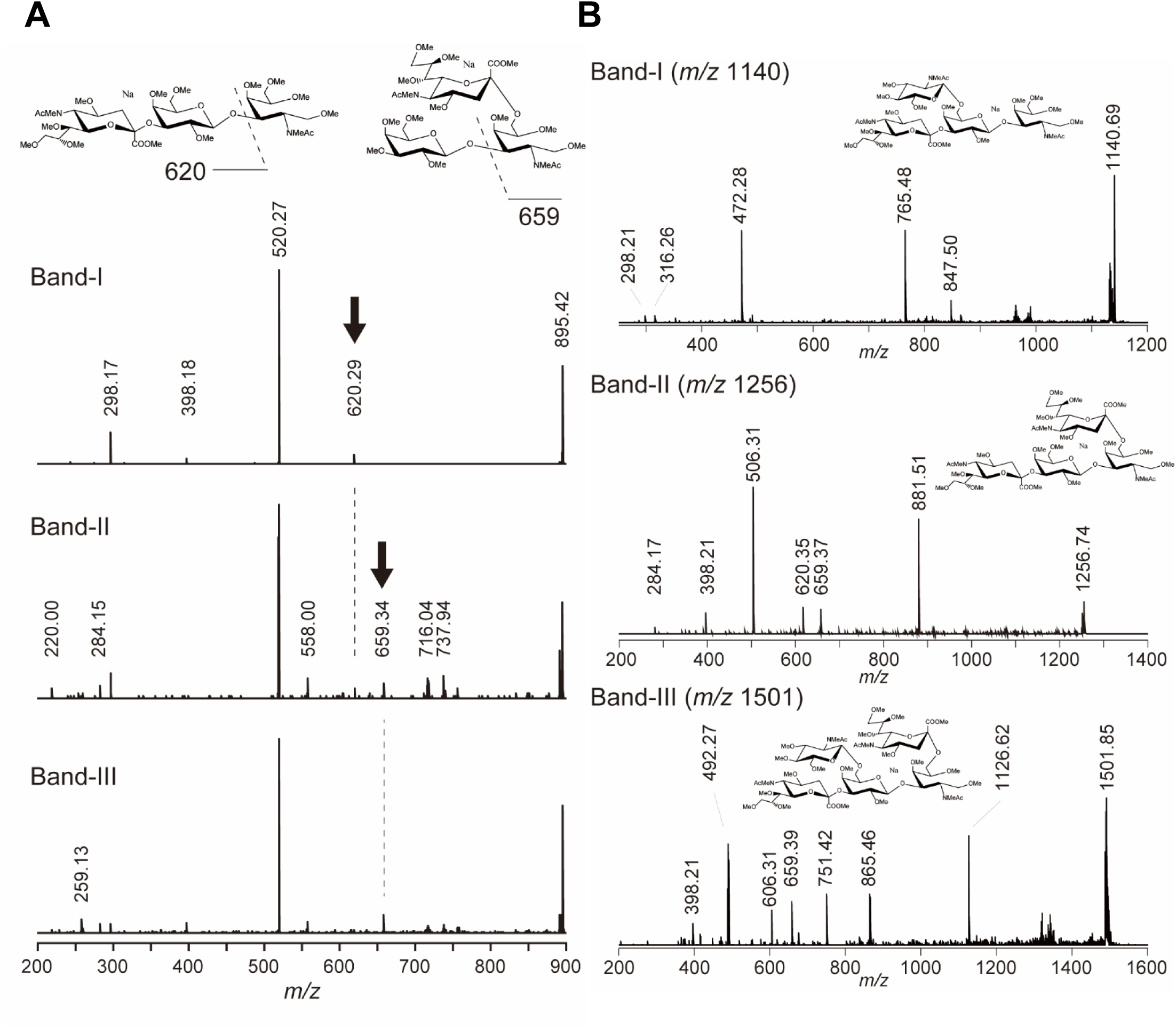
MS/MS spectra and estimated structures of major glycans observed in mouse SMG. (A) MS/MS spectra of glycan 5 (*m/z* 895) in band-I, II, and III. Deduced glycan structures are shown at the top of the spectra. Arrows indicate the key fragments (*m/z* 620 and 659) for the NeuAc2-3Gal and NeuAc2-6GalNAc, respectively. Sample xv was used for MS/MS analysis of glycan 5 in band I and III, and sample i was used for band-II. (B) MS/MS spectra glycan 12 (*m/z* 1140), 15 (*m/z* 1256), and 20 (*m/z* 1501) in band-I, II, and III, respectively. The deduced glycan structure is shown at the top of each spectrum. All spectra were obtained from sample xv. All assignments of fragments in the spectra are shown in eFigure 2.

**Figure 4.**
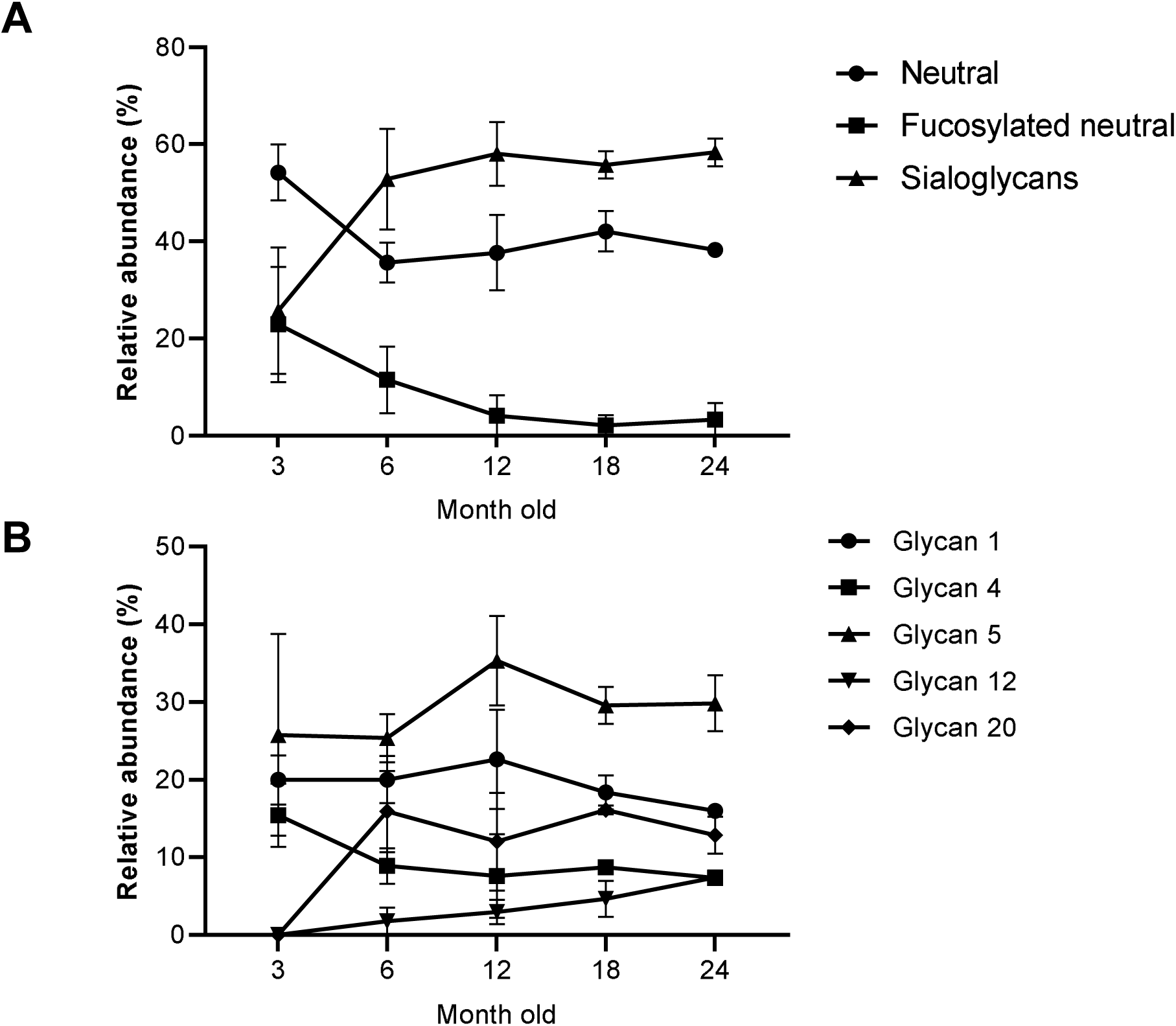
Trends of glycan abundances of band-III during aging. (A) Trends of abundances of three glycan categories: neutral glycans, fucosylated neutral glycans, and sialoglycans. The abundances were the sum of abundance of each glycan in the same category. Data are presented as means ± s.e.; for each age, *n* = 3. (B) Trends of abundancies of major glycans (1, 4, 5, 12, and 20) found in band-III. Data are presented as means ± s.e.; for each age, *n* = 3.

## Results

### Altered SMG mucin production during the aging process in mice

We investigated the reproducibility of expression of the new mucin-like molecule that was specifically detected in aged mice and assessed from which stage of the aging process it was detected. One type of mucin that was not detected in 3-month-old mice appeared with a weak signal at around 12 months of age, gradually became stronger with aging, and was clearly detected again at 24 months of age (Figure 1A, band-I), similar to the mucin-like molecule in a previous report (20). Another type of mucin was abundantly detected in mice of all ages (Figure 1A, band-III), and the third type of mucin was detected only in mice aged 3 to 12 months (Figure 1A, band-II). These results suggest that the mouse SMG has at least three types of mucin (band-I, age-specific mucin; band-II, youth-specific mucin; band-III, mucin common to all ages) with different SMME mobilities. The youth-specific mucin (band-II) was expressed until about 12 months old, and the age-specific mucin (band-I) was expressed after about 12 months old. Mucin common to all ages (band-III) was expressed throughout life. Therefore, it appears that mucin production in the SMG of mice changes with aging.

**Table 1.** Assigned signals to O-glycans in the MS of permethylated alditols obtained from the excised bands.

| Glycan | Obsd. (*m/z*) | Clcd.<sup>a</sup> (*m/z*) | Glycan composition | Classification<sup>b</sup> |
| --- | --- | --- | --- | --- |
| 1 | 534.26 | 534.29 | (Hex)<sub>1</sub>(HexNAc)<sub>1</sub> | N |
| 2 | 708.39 | 708.38 | (Hex)<sub>1</sub>(HexNAc)<sub>1</sub>(dHex)<sub>1</sub> | F |
| 3 | 721.47 | 721.37 | (HexNAc)<sub>1</sub>(NeuGc)<sub>1</sub> | S |
| 4 | 779.46 | 779.41 | (Hex)<sub>1</sub>(HexNAc)<sub>2</sub> | N |
| 5 | 895.53 | 895.46 | (Hex)<sub>1</sub>(HexNAc)<sub>1</sub>(NeuAc)<sub>1</sub> | S |
| 6 | 925.68 | 925.47 | (Hex)<sub>1</sub>(HexNAc)<sub>1</sub>(NeuGc)<sub>1</sub> | S |
| 7 | 953.57 | 953.51 | (Hex)<sub>1</sub>(HexNAc)<sub>2</sub>(dHex)<sub>1</sub> | F |
| 8 | 983.60 | 983.51 | (Hex)<sub>2</sub>(HexNAc)<sub>2</sub> | N |
| 9 | 1024.63 | 1024.54 | (Hex)<sub>1</sub>(HexNAc)<sub>3</sub> | N |
| 10 | 1069.69 | 1069.55 | (Hex)<sub>1</sub>(HexNAc)<sub>1</sub>(dHex)<sub>1</sub>(NeuAc)<sub>1</sub> | S |
| 11 | 1099.70 | 1099.56 | (Hex)<sub>1</sub>(HexNAc)<sub>1</sub>(dHex)<sub>1</sub>(NeuGc)<sub>1</sub> | S |
| 12 | 1140.73 | 1140.59 | (Hex)<sub>1</sub>(HexNAc)<sub>2</sub>(NeuAc)<sub>1</sub> | S |
| 13 | 1157.68 | 1157.61 | (Hex)<sub>2</sub>(HexNAc)<sub>2</sub>(dHex)<sub>1</sub> | F |
| 14 | 1228.72 | 1228.64 | (Hex)<sub>2</sub>(HexNAc)<sub>3</sub> | N |
| 15 | 1256.79 | 1256.64 | (Hex)<sub>1</sub>(HexNAc)<sub>1</sub>(NeuAc)<sub>2</sub> | S |
| 16 | 1286.82 | 1286.65 | (Hex)<sub>1</sub>(HexNAc)<sub>1</sub>(NeuAc)<sub>1</sub>(NeuGc)<sub>1</sub> | S |
| 17 | 1316.83 | 1316.66 | (Hex)<sub>1</sub>(HexNAc)<sub>1</sub>(NeuGc)<sub>2</sub> | S |
| 18 | 1402.97 | 1402.73 | (Hex)<sub>2</sub>(HexNAc)<sub>3</sub>(dHex)<sub>1</sub> | F |
| 19 | 1473.84 | 1473.77 | (Hex)<sub>2</sub>(HexNAc)<sub>4</sub> | N |
| 20 | 1501.85 | 1501.77 | (Hex)<sub>1</sub>(HexNAc)<sub>2</sub>(NeuAc)<sub>2</sub> | S |
| 21 | 1647.84 | 1647.85 | (Hex)<sub>2</sub>(HexNAc)<sub>4</sub>(dHex)<sub>1</sub> | F |
| 22 | 1706.04 | 1705.86 | (Hex)<sub>2</sub>(HexNAc)<sub>2</sub>(NeuAc)<sub>2</sub> | S |
| 23 | 1951.16 | 1950.99 | (Hex)<sub>2</sub>(HexNAc)<sub>3</sub>(NeuAc)<sub>2</sub> | S |

<sup>a</sup> All mass to charge ratios (*m/z*) were calculated as [M + Na]<sup>+</sup> ions of the corresponding permethylated alditols.
<sup>b</sup> N: Neutral glycan; F: fucosylated neutral glycan; S: sialoglycan.

To investigate the relationship between mucin levels and aging, we performed densitometric analysis of the mucin bands detected in Figure 1A. No statistically significant difference was found among the average total band intensities (I + II + III) for each age, but the signal was strongest at 6 months of age and then decreased with age (Figure 1B). Thus, aging may reduce the overall amount of mucins.

### Three types of mucins have different glycan profiles

To examine the relationship between each of the three mucins and their glycan species, we first released and permethylated glycans from mucins at bands I, II, and III and measured them using MS. A total of 23 O-glycans were assigned and summarized in Table 1. Based on monosaccharide composition, the observed glycans could be classified into three groups: neutral glycans, fucosylated neutral glycans, and sialoglycans. The latter two groups consist of fucose (deoxyhexose:deHex) and sialic acid (N-acetyl neuraminic acid: NeuAc; and N-glycolyl neuraminic acid: NeuGc) at the nonreducing glycan ends, respectively, while the former had neither of these. The sialoglycan content decreased from bands I to II to III, while that of fucosylated neutral glycan increased (Figure 2). Most of the glycans of band-I, the age-specific mucin, were sialoglycans (Figure 2, band-I). Glycan 5 (*m/z* 895) was the most abundant in all three bands, while the second peak was characteristic of each band: glycan 12 (*m/z* 1140) for band-I, glycan 15 (*m/z* 1256) for band-II, and glycan 20 (*m/z* 1501) for band-III (Figure 2), indicating that the three types of mucins contained different glycan profiles.

### The primary glycan in all mucins, glycan 5 (*m/z* 895), has different structures on each mucin

We next estimated the detailed structure of the observed O-glycans using tandem MS. Interestingly, the results revealed that the glycan 5 at *m/z* 895, which is the most abundant glycan in all three bands I, II, and III, has different glycan structures in each band (Figure 3A). Tandem MS of glycan 5 (NeuAc-Hex-HexNAc) afforded characteristic fragments *m/z* 620 and 659, which are exclusively derived from the NeuAcα2-3Gal and NeuAcα2-6GalNAc structures, respectively. In the MS/MS spectra of glycan 5 from bands I and III, the signature fragments were detected at *m/z* 620 and 659, respectively, indicating that the glycan structures of band-I and band-III are NeuAcα2-3Galβ1-3GalNAc and Galβ1-3(NeuAcα2-6)GalNAc, respectively. Band-II afforded both fragments, indicating it was a mixture of the two structures (Figure 3A). In addition, SMME membrane blots with MAL-II lectin that bind to core 1 type NeuAcα2-3Gal structure stained band-I and II (eFigure 1). This also supported that the major glycans of band-I and II mucins had a structure of NeuAcα2-3Galβ1-3GalNAc. Next, the glycans 12, 15, and 20, which are the second or third major glycans of bands-I, II, and III, were estimated using tandem MS as follows: the *m/z* 1140: NeuAcα2-3 (GlcNAcβ1-6)Galβ1-3GalNAc, *m/z* 1256, NeuAcα2-3Galβ1-3(NeuAcα2-6)GalNAc, and *m/z* 1501, and NeuAcα2-3(GlcNAcβ1-6)Galβ1-3(NeuAcα2-6)GalNAc, respectively (Figure 3B). Assignments of fragments in the MS/MS spectra are summarized in eFigure 2. The structures of glycan 3, 4, and 6 were also estimated. The monosaccharide composition of glycan 4 is (Hex)_1_(HexNAc)_2_ that invokes core-2 structure for O-glycans, Galβ1-3(GlcNAcβ1-6)GalNAc. However, tandem MS revealed that glycan 4 has a linear structure GlcNAc-Gal-GalNAc but not core-2. Given the biosynthetic pathway of O-glycans and the estimated structures for other glycans observed in this experiment, we estimated that glycan 4 was GlcNAcβ1-6Galβ1-3GalNAc (eFigure 3A). The structures of glycans 3 and 6 containing NeuGc (N-glycolylneuraminic acid) were also confirmed using tandem MS (eFigure 3B). All mucins of bands I, II, and III have a small proportion of glycan 3 and glycan 6. This suggests that the mouse SMG contains CMP-NeuAc hydroxylase (24,25). It became clear from these data that the glycan species on each of the three mucins and their proportions are different. If the mucin core protein is different, it is highly likely that the glycan species is different (Robbe, Capon, Coddeville, & Michalski, 2004). Therefore, it is speculated that the core proteins of the three mucins will be different.

### The glycans of band-III, a mucin common to all ages, change with aging

To investigate the effect of aging on the glycan species of band-III, a mucin common to all ages, percentages of particular glycan species in band-III were plotted every 3 months. Between 3 and 6 months of age, the proportion of fucosylated neutral glycans and neutral glycans decreased, while that of sialoglycans significantly increased (Figure 4A). After 6 months of age, neutral glycans and sialoglycans remained almost constant, while the ratio of fucosylated neutral glycans decreased during the aging process (Figure 4A). The proportion of major glycans 1, 4, and 5 did not change significantly, but that of sialoglycan 20 increased sharply between 3 and 6 months and that of sialoglycan 12 increased with aging (Figure 4B). These results suggest that for SMG mucin common to all ages (band-III), aging tends to decrease the proportion of fucosylated glycans and increase the proportion of sialoglycans.

## Discussion

Maintaining oral function is extremely important for the elderly to live a healthy and high-quality life, but many elderly people have xerostomia with symptoms such as decreased salivary secretion and increased salivary viscosity. Xerostomia in the elderly not only causes various disorders such as dental caries and periodontal disease (Abdellatif & Burt, 1987; Gilbert et al., 1993), denture incompatibility (Locker, 1993), and oral mucosal disease (Epstein, Pearsall, & Truelove, 1980; Hopcraft & Tan, 2010; Osterberg et al., 1984; Sreebny, 2000) but also affects the whole body as with the increased risk of aspiration pneumonia (Sjogren et al., 2008; van der Maarel-Wierink et al., 2013). Therefore, xerostomia significantly impacts the quality of life of otherwise healthy elderly people.

Mucins, produced in the SGs, have large masses and complex biochemical compositions consisting of 50%–80% glycans, and tend to form higher-order structures due to polymerization (Wu et al., 1994). For this reason, the molecular science on mucins is lagging, and studies on xerostomia rarely focus on mucins in SGs. Using SMME, a method developed to isolate high molecular weight glycoproteins (Matsuno et al., 2009), we have previously studied both membrane-bound and secreted mucins in the SMG of mice (Iida et al., 2019) and have reported that aging leads to the expression of a new secreted mucin-like molecule (Iida et al., 2019; Kameyama et al., 2019). Therefore, in the present study, to investigate in more detail the secretory mucin produced and secreted in the SMGs during the aging process in mice, we slightly modified a mucin enrichment method to perform mucin separation and subsequent mass spectrometry to analyze the glycan structure on mucin molecules from the SMG. We thus discovered three secreted mucins with different glycan profiles in the SMGs of mice (age-specific mucin band-I, youth-specific mucin band-II, and mucin band-III that is expressed throughout of life), and that each mucin has its own peak production age during the aging process. Consistent with our previous findings, age-specific mucin was a sialomucin containing mostly sialoglycans. In all three mucins, the main glycan 5 at *m/z* 895, had the same mass among mucins, but the glycan structure was different for each mucin. The total amount of mucins, including these three acidic mucins, reached a maximum at 6 months of age, and thereafter decreased with aging. This is consistent with a report examining the effect of aging on mucin content in the mouse SMG (Denny, Klauser, Villa, Hong, & Denny, 1991). In our previous study, we also attempted to identify mucins detected by SMME using various analytical methods and found that the mucins expressed throughout life are Prol1/Muc10 (Kameyama et al., 2019). However, details of the age-specific mucin are still unknown. Several previous studies on mucins have shown that different core proteins have different glycan species (Robbe et al., 2004), and that identical core structures do not always result in the same glycosylation pattern and are affected by a number of additional factors such as tissue and cell type, substrate availability, and physiological environment (Carraway & Hull, 1989; Derrien et al., 2010; Lan, Batra, Qi, Metzgar, & Hollingsworth, 1990). In the present analyses of glycan species in SMG mucins, we found that each of the three mucins had different glycan species and proportions. Taken together with previous studies, these data suggest that the core proteins of the mucins are different.

This may imply that more than one type of cell produces mucins in the SMG. However, there is one type of acinar cell that secretes both serous and mucous components in the mouse SMG (Amano, Mizobe, Bando, & Sakiyama, 2012). In addition, the SMG of mice has a granular duct (granular convoluted duct, GCT) that produces and secretes various bioactive polypeptides, hormones, and cell growth factors, but these cells do not have sialoglycoconjugates (Accili, Gabrielli, Menghi, & Materazzi, 1996). Furthermore, GCT cells are not stained by AB in young or aged mice (Iida et al., 2019). Therefore, it is unlikely that the type of cells producing this mucin increases due to aging. Thus, we propose that the three mucins detected by SMME analysis are produced in acinar cells and composed of different core proteins. Next, we found that the percentage of mucin glycan species expressed throughout life changes during the aging process, and aging decreases the proportion of fucosylated neutral glycans and increases the proportion of sialoglycans. Therefore, it is presumed that changes in the physiological environment due to aging fluctuates the proportions of each glycan species of mucin expressed throughout life.

Salivary mucins are components of the tooth pellicle and play a role in colonizing oral bacteria on the tooth pellicle. Recent studies have shown that *Streptococcus gordonii*, one of the oral streptococci that colonizes the tooth surface, binds to host sialic acid (Blix, Svennerholm, & Werner, 1952; Schauer, 1982) via sialic acid-binding adhesin (Takahashi, Sandberg, Ruhl, Muller, & Cisar, 1997). Sialomucin also contains a significant amount of negatively charged sialic acid, which gives the mucin molecule a strong negative charge (Derrien et al., 2010). This study and our previous studies have shown that aging results in the expression of age-specific sialomucin (Iida et al., 2019; Kameyama et al., 2019) as well as an increase in the ratio of sialoglycans in the SMG mucin expressed throughout life. Therefore, aging likely causes many mucin molecules to have a strong negative charge. Sialoglycans, which increase with aging, and the negative charge that they bring to mucin molecules therefore affect mucin polymerization and the ability to bind various microorganisms. In this way, they putatively alter mucin-facilitated microbial clearance, microbial colonization and colony formation, and mucin degradation by microorganisms. Therefore, it is suggested that the cause of the increase in salivary viscosity and oral hygiene deterioration that occurs with age is likely to be such changes in mucin functions in addition to a decrease in saliva flow and water evaporation (Dawes, 1987; Narhi, 1994).

This study revealed that in the mouse SMG, mucin production and glycoforms are altered during aging. Interestingly, inflammation has been reported to induce mucin overexpression, inappropriate expression, or abnormal forms of expression (Rose & Voynow, 2006). Aging causes inflammation in various tissues. This is also the case with the SGs, and in the SGs of aged mice, the infiltration of inflammatory cells increases, and inflammation occurs (Choi, Park, Kim, Lim, & Kim, 2013). Thus, the inflammatory response generated by aging may alter the production and glycoforms of mucin in the SMGs.

Herein, we focused on mucins in the SMGs secreting saliva and examined the nature of SMG mucin during the aging process. We found that mouse SMGs produce three types of secreted mucins with different glycan profiles and peak production ages. In one of these mucins, expressed throughout life, different proportions of glycan species are expressed during the aging process. This suggests that changes in the physiochemical structure and function of mucin may contribute to the increase in salivary viscosity and deterioration of oral hygiene that are observed during the process of aging. However, the process of aging influences many factors such as muscle weakness, diabetes, increased medication, and stress, and these combined effects produce xerostomia (Bergdahl & Bergdahl, 2000; Dusek, Simmons, Buschang, & al-Hashimi, 1996; Lopez-Pintor, Casanas, & Gonzalez-Serrano, 2016; Murray Thomson, Chalmers, John Spencer, Slade, & Carter, 2006). Therefore, in elucidating the causes of xerostomia, it is necessary to understand the physicochemical structures and functions of mucins contained in the saliva of the elderly while also accounting for the relationship with these factors.

## Acknowledgments

We thank the Laboratory of Experimental Animals of the National Center for Geriatrics and Gerontology for providing the aged mice.

## Appendix A. Supplementary data

The following is Supplementary data to this article: Supplementary Figure 1, Supplementary Figure 2, Supplementary Figure 3.

